# Fitness effects of copper selection in a wild-derived population of *Drosophila melanogaster*

**DOI:** 10.64898/2026.08.20.746007

**Authors:** Christina M Rodriguez, Kazzrie A Arnold, Elizabeth R Everman

## Abstract

Copper is an essential micronutrient in most organisms that becomes toxic in large quantities. Repeated or prolonged sub-lethal exposure can lead to evolved resistance to copper toxicity over many generations, which may result in trade-offs between energetically expensive detoxification mechanisms and fitness. Alternatively, evolved resistance to chemical stressors may lead to correlated changes in other traits. This study focuses on a population of flies for which artificial selection for copper resistance led to an increase in both copper resistance and longevity. The apparent off-target benefit of copper selection on one component of fitness led us to investigate differences in fecundity and developmental viability in copper resistant and copper sensitive, non-selected populations. We assessed the effect of copper selection and copper exposure on multiple aspects of fecundity over the lifespans of females from the non-selected and copper-selected populations. Our study corroborated previously observed increased longevity in copper-selected flies. Controlling for variation in lifespan, copper-resistant females had comparable age-matched fecundity to copper-sensitive females and benefitted from increased longevity with higher lifetime fecundity. Overall, copper exposure negatively affected egg quality, but we found no difference in this trait between the copper-resistant and sensitive populations. Further, we found developmental viability under copper stress was significantly higher for eggs laid by copper-resistant females. Overall, we determined that copper resistant flies experienced a fitness benefit through both lifespan and fecundity. Costs of maintaining copper resistance may be associated with energetic costs, but these trade-offs may not always manifest in reproductive or lifespan fitness costs.

## Introduction

Adaptation to physiological stressors has the potential to influence traits that are not the direct target of selection due to overlap in genetic control or to physiological, biochemical, functional, or energetic constraints (Stephen 1992; Garland et al. 2022). These correlated responses are often predicted to result in trade-offs with fitness, especially when optimization of resistance is achieved at the expense of resources for growth and/or reproduction (ffrench-Constant and Bass 2017). Trade-offs between chemical stressors and fitness have been observed in several organisms including insecticide resistance in mosquitoes (*Culex pipiens*), heavy metal resistance in worms (*Neris diversicolor*), and heavy metal tolerance in hyperaccumulating plants (Küpper et al. 2001; Pook et al. 2009; Maestri et al. 2010; Rivero et al. 2011). In some cases, trade-offs are driven by energetic or metabolic costs that deplete lipid, glycogen, and glucose reserves and are associated with reduced growth rate and fecundity (Pook et al. 2009; Rivero et al. 2011). In other cases, such as for many hyperaccumulating plants, consequences for fitness traits may be more complex, with some plant species displaying a ‘typical’ trade-off between heavy metal resistance and growth rate and others relying on high metal concentrations for their own optimal growth and reproduction (Küpper et al. 2001; Maestri et al. 2010).

The benefits and consequences of adaptation to heavy metal contamination may be more complex for heavy metals that function as essential micronutrients. Resistance to high environmental concentrations of copper or zinc, for example, cannot be achieved by simply inactivating uptake of these essential metals. Instead, organisms must adapt to the increased influx of ions through modification of complex detoxification pathways. Copper is one such essential heavy metal that is also a common environmental contaminant released through agricultural and industrial pollution. Overexposure results in oxidative stress-induced toxicity that can damage lipids, DNA, proteins, and tissues over short term exposure (Husak 2015; Calap-Quintana et al. 2017). Accumulation of damage can ultimately lead to organ failure and death under acute or prolonged stress conditions (Husak 2015). Even if exposure concentrations are sublethal, elevated copper has the potential to impact multiple aspects of an organism’s physiology and reproduction. For example in aquatic invertebrates (Cladocera), chronic exposure to copper reduced growth rate, reproduction, and fecundity, and negatively affected development (Sadeq and Beckerman 2019).

While negative effects of metal exposure on physiology and reproduction are fairly common in a wide range of organisms (Georgieva et al. 2002; Jomova et al. 2010; Das et al. 2017), populations may adapt over time to consistent exposure to metal stress (Eränen 2008; Gerstein et al. 2015; Bazzicalupo et al. 2019; Papadopulos et al. 2021; Everman et al. 2023; Arnold and Everman 2026). This can occur through adaptation, acclimation or a combination of these mechanisms. When the effects of acclimation and adaptation are separated, anticipated trade-offs are not always observed. For example, in earthworms (*Dendrobaena ocaedra*) that inhabit copper-contaminated soils (contaminated for 300 years with industrial pollution), copper-adapted individuals had faster individual growth rates compared to non-adapted individuals, suggesting negligeable costs of copper adaptation (Fisker et al. 2011). Similarly in *Daphnia longispina* collected from an abandoned mine drainage site, copper-adapted strains had faster population growth rates compared to sensitive strains (Agra et al. 2011). However, in a recent study of copper adaptation using artificial selection in *Drosophila melanogaster*, we found that as populations that were subjected to selection gained copper resistance, control populations (not subjected to selection) lost resistance to copper, cadmium, and lead over many generations (Arnold and Everman 2026). The loss of heavy metal resistance in control populations suggests that there is a cost of maintaining resistance, which may stem from energetically expensive detoxification processes requiring gene upregulation and increased metabolic activity to bind excess metal ions, detoxify, and excrete them (Calap-Quintana et al. 2017; Ruiz et al. 2021; Osborne and Prosser 2025a).

Arnold and Everman (2026) investigated the possible costs of copper resistance by measuring longevity of copper-selected and control populations, and found that copper-selected female flies outlived control females by 13 days under control conditions. Several prior studies in *D. melanogaster* have reported a similar positive association between stress resistance and longevity (particularly between longevity, starvation, and oxidative stress resistance (Service et al. 1985; Harshman et al. 1999)). However, trade-offs between heavy metal resistance and fitness traits have also been reported. Notably, Shirley and Sibly (1999) found a clear trade-off between cadmium resistance and fecundity in flies that had experienced 20 generations of artificial selection for resistance to cadmium stress. While the cadmium-resistant flies had higher survival and fecundity on cadmium-contaminated media compared to sensitive flies, resistant flies lost these advantages on control media. Given these prior observations of a metal resistance-fecundity trade-off, we investigated potential fitness trade-offs with copper resistance by assessing age-specific and total fecundity, egg quality, and developmental success of the offspring of copper-selected and copper-sensitive non-selected flies under control and copper stress conditions.

## Methods

### Fly culturing

The flies used in this study were descended from wild-caught females originally collected from a commercial orchard in Blue Ridge, Georgia (USA) in July 2024. Following collection, the offspring of each wild female were allowed to mate freely in cages as outbred populations (BugDorm, 20cm x 20cm x 20cm) as described in detail in (Arnold and Everman 2026). Briefly, one set of population cages were maintained under consistent control conditions with control (non-copper) food. The other set of population cages were subjected to selection for copper resistance in the adult life stage every other generation starting with generation five. Every other generation, adult flies in the copper selected cages were given sole access to media containing 50mM Copper(II) Sulfate (50mM CuSO_4_; Sigma-Aldrich C1297) until approximately 50% of the population died. Eggs were collected from surviving females to establish the next generation. The control (MC) and copper-selected (MA) population cages that are the subject of the current study are part of a larger, on-going artificial selection experiment. We chose to focus on these two cages because the copper-selected population evolved increased lifespan in addition to increased copper resistance (see Figure 5 of Arnold and Everman (2026)). Control and copper-selected cages and all experiments detailed below were housed in Percival *Drosophila* Incubation Chambers at 25°C and 50% relative humidity with a 12-hour light:dark cycle.

### Collection of eggs for phenotyping

To assess all traits, eggs were collected from the MC and MA cages using apple juice plates (apple juice, cane sugar, 1% agar, and propionic, phosphoric, and benzoic acid media preservatives) baited with a small daub of yeast paste (equal volume dry active yeast and ePure water). When flies in each cage were approximately 3-5 days old, one apple juice plate was placed in each cage for 24 hours. Eggs were collected by suspending them in 1x PBS and pipetting 30uL of eggs into vials containing standard *Drosophila* media (recipe available in Everman et al. (2019)). When flies were between 3 days old, males and females were sorted with light CO_2_ anesthesia into same-sex groups for each experiment and were transferred to experimental vials after 24 hours of recovery from sorting.

### Adult lifespan

Flies were sorted into same sex groups of 10 individuals per vial (N = 49-50 replicate vials/sex/population cage) to estimate adult lifespan. Fewer flies were included in each vial (in comparison to other experiments described below) to facilitate accurate manual counts of live flies. Survival was assessed every Monday, Wednesday, and Friday of the experiment until all flies were dead. Flies were transferred to new food (without copper) on each day they were counted to minimize the impact of vial conditions on the survival of flies. We calculated average lifespan per vial replicate as the response variable.

### Adult copper resistance

Flies were sorted into same sex groups of 20 individuals per vial (N = 5 replicate vials/sex/population cage) for estimation of adult copper resistance. Adult copper resistance was assessed by transferring flies without anesthesia to vials containing the same copper concentration used for selection (50mM CuSO_4_). Copper treatment was administered by hydrating 1.8g Instant Drosophila Media (Carolina Biological Supply Company 173200) with 8mL 50mM CuSO_4_. Survival was assessed daily by counting dead flies until all flies were dead. We calculated average lifespan per vial replicate as the response variable.

### Fecundity

Age-specific and lifetime fecundity were estimated in mated females beginning with 3-day old flies. Females (N = 5 individuals/vial) were housed with age-matched males from the same population cage as the females (N = 5 individuals/vial) at the beginning of the experiment. Males were not replaced. We assessed fecundity of non-selected and copper-selected females under control and copper-stress conditions using a fully factorial design (N = 10 vials/selection history/copper stress treatment). To ensure that copper-stressed females survived to an old age that was not drastically different from control conditions, we used a lower concentration determined from pilot experiments (5mM CuSO_4_) compared to the adult copper resistance assay described above. All flies were provided with high protein media (Backhaus et al. 1984), with or without copper, through the duration of the experiment to encourage egg laying (Lee et al. 2008).

To assess fecundity, we used removable plastic caps (MOCAP FCS13/16NA1) that could be easily photographed and replaced in narrow *Drosophila* vials (Genessee Scientific 32-120). Each cap contained approximately 5mL of copper or control media and was labeled according to vial replicate and treatment. Females were allowed 24 hours to lay eggs on caps, after which the caps were photographed using a DSLR camera (Nikon D780, SIGMA 105mm 1:2.8 DG MACRO HSM) and replaced. We recorded the number of surviving females each Monday, and eggs on cap photographs were manually counted using the Cell Counter plugin, part of ImageJ (Abràmoff et al. 2004; Rasband 2007) from Monday – Wednesday, providing three replicate estimates of fecundity per week of the experiment. Each Thursday, flies were transferred to semi-defined media according to control or copper treatment in vials until the following Monday, and all transfers were done without anesthesia. The experiment continued until all females were dead.

To account for the decline in number of females in each vial over the course of the experiment, the number of eggs per cap was adjusted by dividing the total eggs laid by the number of females alive, providing an average estimate of eggs laid per female for each vial replicate as the response variable. In addition, we estimated average lifetime fecundity per female by summing the number of eggs (corrected for number of females alive at the start of the week) laid in each cap. As we only counted eggs laid for three days of each week, our estimates of average lifetime fecundity are likely conservative.

### Egg length

We used estimates of egg length to assess changes in egg quality over the duration of the fecundity experiment. Using the same images that were used to estimate fecundity, we measured the length of up to 10 eggs on each cap on the first fecundity assessment of each week. To control for variation across caps and across images, we calibrated the mean length of pixels for each individual cap using the Line tool in ImageJ. We then estimated the length of each egg in pixels with the Line tool and converted values to millimeters using the known diameter of the cap and the “set scale” option in ImageJ. Egg length was only estimated for eggs that were laying entirely flat against the surface of the media, and spicules were not included in estimates. Average egg length was calculated for each cap and was used as the response variable (N = 10 caps/treatment/population cage/week).

### Developmental success

To assess viability of eggs laid by females from the copper-selected and non-selected cages, we collected eggs from the cages using apple juice plates baited with yeast paste. Parents of eggs (approx. 3 days old) were not exposed to copper prior to egg collection. We manually picked sets of 50 eggs from the apple juice plates and placed them into vials containing the same high-protein media used to assess fecundity containing no copper (control) or 1mM CuSO_4_, 5mM CuSO_4_, or 7mM CuSO_4_ (N = 3 vials/treatment/population cage). We chose a limited range of copper concentrations including the concentration used in fecundity assays to explore copper effects on developmental success under low and moderate stress conditions (concentrations were chosen using pilot experiments, data not shown).

Experimental vials were monitored daily, and the number of eclosing male and female flies was recorded once per day 3 hours after lights turned on in the incubators. For each experimental vial, we calculated sex-specific and total viability (proportion of eggs that emerged as adults), average time (days) to emergence, and sex ratio (males to females).

### Data analysis

For traits with a single response (adult lifespan, adult copper resistance, lifetime fecundity, developmental viability, time, and sex ratio), we used separate analyses of variance (ANOVA, type III sums of squares) using the base R ‘lm’ function and ‘Anova’ from the car package (Fox et al. 2001; R Core Team 2023). We tested the additive and interaction effects of Sex and Selection History on adult lifespan and copper resistance with two-way ANOVAs. After an initial assessment of Sex, Selection History, and Copper Treatment with a three-way ANOVA, we found that Sex did not significantly influence development viability (F_(1,32)_ = 0.54, P = 0.46) or development time (F_(1,31)_ = 0.84, P = 0.37). Therefore, we proceeded with two-way ANOVAs for these traits and sex ratio to estimate the additive and interaction effects of Selection History and Copper Treatment. Parametric assumptions were tested using the Shapiro-Wilk test (base R) for normality and the Levene test (car package) for homoscedasticity.

No single response traits deviated from parametric assumptions except for lifetime fecundity, development time, and sex ratio. Lifetime fecundity deviation from normality did not improve with data transformation, and because the data were homoscedastic (F_(3,26)_ = 0.50, P = 0.23), we proceeded with a parametric test of untransformed data. Development time and sex ratio were both right-skewed but homoscedastic as well (Development Time: W = 0.83, P < 0.002, Levene test: F_(7,16)_ = 1.44, P = 0.26; Sex Ratio: W = 0.78, P < 0.0002, Levene test: F_(7,16)_ = 2.15, P = 0.1). Deviations from normality were improved by square-root transforming both responses, so the two-way ANOVAs were performed on transformed data for development time and sex ratio.

When significant, the main and interaction effects were investigated further using the function ‘emmeans’ to compare population cages with different selection histories within copper treatment, maintaining an experiment-wide alpha level of 0.05 (Bonferroni correction) (Lenth 2025).

Fecundity and egg length were both assessed throughout the duration of the experiment and were analyzed with analyses of covariance (ANCOVA). In both analyses, we used the proportion of surviving females as the covariate instead of chronological time to account for differences in longevity due to copper treatment or due to other unmeasured innate differences between the two populations. The main effects and interactions between Copper Treatment, Selection History, and Female Survival were included in the model. Because the corrected number of eggs laid per female was right-skewed, these data were square-root transformed, which improved violation of normality and variance assumptions (W = 0.99, P < 0.00003; Levene test: F_(3,851)_ = 3.49, P < 0.02). Transformation of egg length did not improve violation of normality assumptions, but because the violation was minor (W = 0.99, P < 0.03), likely influenced by large sample size, and data were homoscedastic (F_(3,239)_ = 2.5, P = 0.06), we proceeded with parametric analysis of raw data.

Data visualization was achieved using the rmisc and tidyverse packages (Hope 2012; Wickham et al. 2019). An alpha level of 0.05 was used to assess significance for all tests, correcting for post-hoc comparisons as appropriate.

### Data availability

All phenotype data generated in this study will be available from Dryad (DOI: https://doi.org/10.5061/dryad.7pvmcvf9h).

## Results

### Copper-selected flies outlived non-selected flies under nonstress and copper-stress conditions

Arnold and Everman (2026) reported that selection for copper resistance in outbred, wild-derived populations of flies collected from a commercial orchard also resulted in a significant increase in longevity. Selection for adult copper resistance has continued nearly every other generation since the initial finding was reported. Consistent with the initial observation, flies (averaged across sex) with a copper-selection history lived nearly six days longer than non-selected flies (Selection History: F_(1,195)_ = 41.54, P < 0.00001) (**Figure 1**A). Females outlived males regardless of selection history (Sex: F_(1,195)_ = 67.60, P < 0.00001; Sex x Selection History: F_(1,195)_ = 3.70, P = 0.06) (**Figure 1**A).

**Figure 1.**
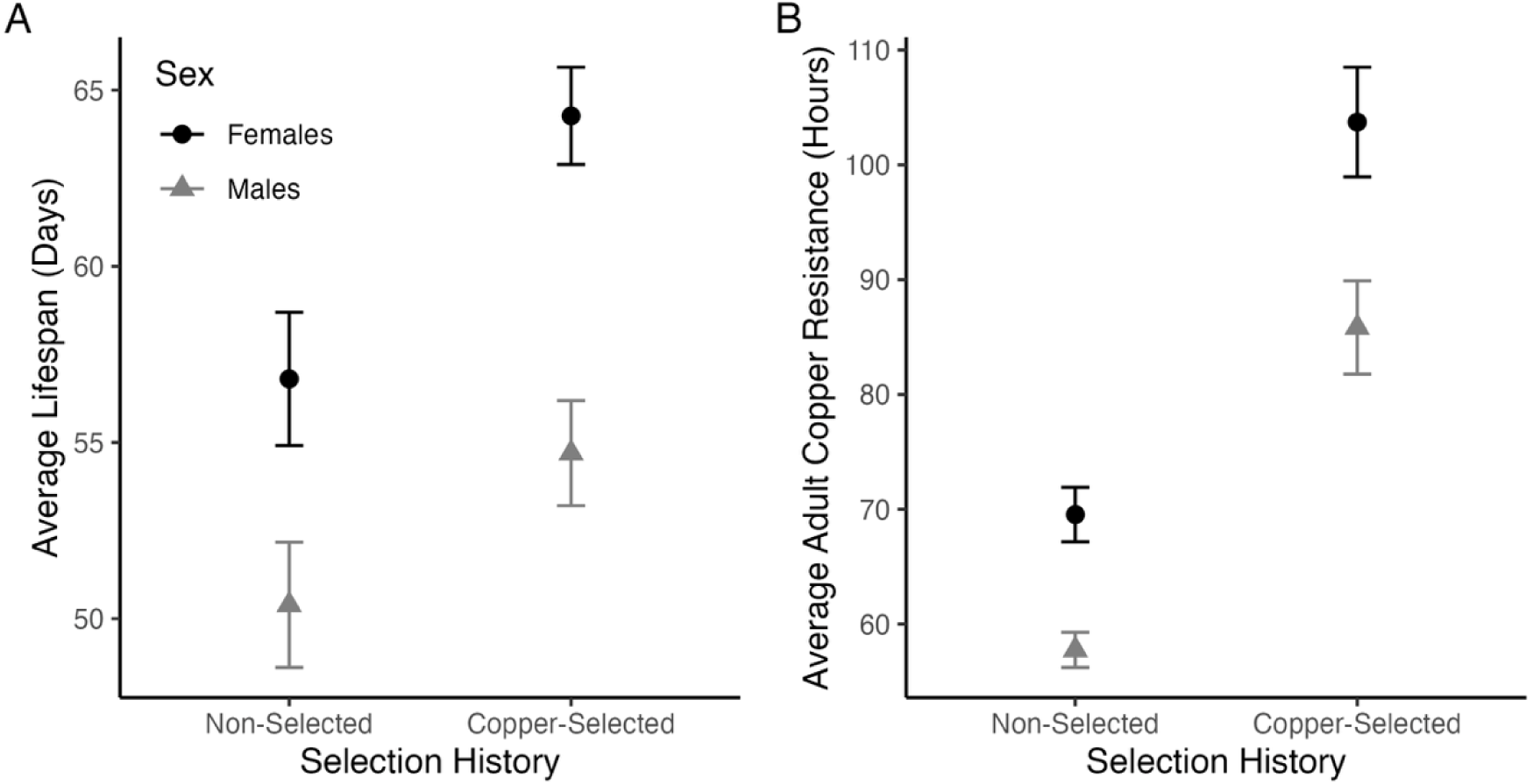
Copper selection increased lifespan and adult copper stress resistance. A. Male and female flies from population cages subjected to artificial selection for copper resistance outlived flies from non-selected populations under non-stressful conditions (F_(1,195)_ = 41.54, P < 0.00001). B. Copper resistance continued to increase in selected populations relative to previous reported resistance in Arnold and Everman (2026), with copper-selected flies living approximately 31 hours longer than non-selected flies when exposed to 50mM CuSO_4_ (F_(1,56)_ = 227.34, P < 0.00001). In both plots, mean estimates +/− 95% confidence intervals are shown. Dark circles indicate average female responses; grey triangles indicate males in both plots. **Alt text:** Graphs showing average lifespan in days and adult copper resistance in hours of male and female flies in two panels labeled A and B. Data shown are summary statistics that illustrate differences between sexes and populations with different selection histories

Copper-selected and non-selected populations have continued to diverge phenotypically, with the resistant flies living on average 31 hours longer than the non-selected flies (Selection History: F_(1,56)_ = 227.34, P < 0.00001) (**Figure 1**B). The difference in resistance was influenced both by a slight continued increase in copper resistance in the selected flies and a decrease in resistance in the non-selected cages (data not shown, see Figure 1 of Arnold and Everman (2026)). Copper resistance was influenced by sex, with females consistently surviving copper stress longer regardless of selection history (Sex: F_(1,56)_ = 62.27, P < 0.00001; Sex x Selection History: F_(1,56)_ = 3.62, P = 0.06). Throughout the process of selection for adult copper resistance, copper-selected cages and control (non-selected) cages were maintained on the same schedule to ensure that all flies in the parent generation were within the same age range regardless of selection regime. As a result, it is unlikely that population maintenance contributed to the increase in longevity observed in the population subjected to copper selection.

### Copper-selected and non-selected females had comparable fecundity and egg quality

Fecundity was estimated by counting the number of eggs laid by females each week until all females were dead. The number of eggs per observation was corrected by the number of females alive to generate per-individual estimates. Because the copper-selected and non-selected females had different lifespans (**Figure 1**), we used female survival at the start of each week as a proxy for the physiological age to test the effects of copper exposure (direct exposure of females used in the fecundity experiment) and selection history on the number of eggs laid over time.

Female age (proxied by survival) significantly influenced fecundity, with females laying fewer eggs as they approached the end of their life (F_(1,847)_ = 452.1, P < 0.0001; **Figure 2**A). Exposure to 5mM CuSO_4_ throughout the duration of the fecundity experiment also had a significant negative effect on fecundity (F_(1,847)_ = 256.7, P < 0.0001) regardless of female age (Treatment x Female Survival: F_(1,847)_ = 2.25, P = 0.13). Similar to longevity, females benefitted from selection for copper resistance by having slightly higher fecundity than non-selected females (F_(1,847)_ = 7.86, P < 0.006; **Figure 2**A and B). A significant interaction between selection history and female survival (F_(1,847)_ = 4.47, P < 0.04) suggests that longer-lived flies may have lower early-life fecundity compared to non-selected females, which is consistent with classic studies in life history that have previously investigated the relationship between longevity and reproduction (Rose 1984; Rose et al. 1992; Zwaan et al. 1995). However, this interaction is minor and is not apparent when fecundity is compared in chronological time (copper-selected females lay the same number of eggs in their first week of the experiment compared with non-selected females; data not shown).

**Figure 2.**
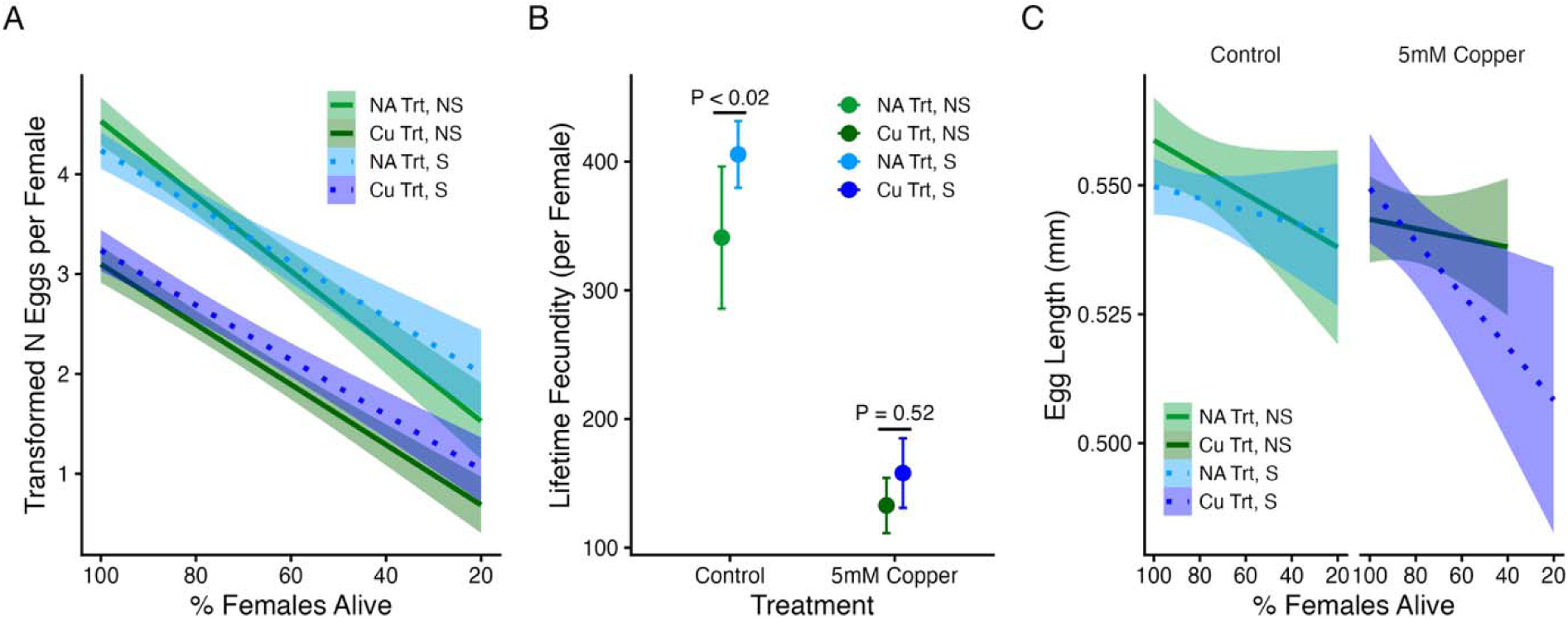
Fecundity and egg quality were influenced by copper treatment or selection history. A. As females aged, they laid fewer eggs and copper treatment had a negative effect on fecundity overall. B. Lifetime fecundity was negatively affected by concurrent copper treatment, but females with a selection history for copper resistance had higher fecundity overall compared to non-selected females. C. Egg length declined with physiological age and in response to copper treatment; however, selection history had a very minor effect only under copper stress conditions (three-way interaction F_(1,235)_ = 4.0, P = 0.047). In all plots, NS (green) indicates females with no selection history for copper stress; S (blue) indicates females with a selection history for adult copper resistance. In A and C, selection history is also indicated by line type (dashed lines indicate copper-selected females). Lighter colors indicate females assessed under control conditions; darker colors indicate exposure to 5mM CuSO_4_ throughout the duration of the experiment. In A and C, the line represents the best fit line of the regression and shading indicates 95% CIs. In B, points indicate estimated mean responses +/− 95% CI. **Alt text:** Graphs showing the change in fecundity in response to selection history and experimental treatment conditions. Subfigures are labelled A through C and show data summaries and statistical results for the number of eggs laid per female over time, lifetime fecundity, and the length of eggs, respectively. Subfigure B provides post hoc analysis of differences in lifetime fecundity due to experimental and selection conditions.

In contrast to our expectation that there would be a trade-off between longevity and fecundity that is more pronounced in copper-selected flies, we found that our long-lived, copper-selected females had higher lifetime fecundity than non-selected females overall and under control conditions (Selection History: F_(1,36)_ = 8.68, P < 0.006; Bonferroni adjusted post hoc test: P < 0.02; **Figure 2**B). Exposure to 5mM CuSO_4_ had a negative effect on overall fecundity (Copper Treatment: F_(1,36)_ = 90.2, P < 0.0001); however, the effect of selection history on fecundity not significant under copper stress (Bonferroni adjusted P = 0.5; **Figure 2**B).

Lastly, we assessed the effects of copper treatment and selection history on egg length to test whether the benefit of increased longevity and apparent lack of cost to fecundity was countered by a trade-off in egg quality. Both the physiological age of females and exposure to 5mM CuSO_4_ negatively affected egg length (Survival: F_(1,235)_ = 10.22, P < 0.002; Copper Treatment: F_(1,235)_ = 8.2, P < 0.005; **Figure 2**C). Although we did not observe an effect of selection history (Selection History: F_(1,235)_ = 1.9, P = 0.17), we did find a weak interaction between selection history, female physiological age, and copper treatment (three-way interaction: F_(1,235)_ = 4.0, P = 0.047) that was primarily influenced by a more pronounced decline in egg length in selected flies under copper stress conditions (**Figure 2**C). This minor effect of copper selection on egg quality under stressful conditions was the only weak indication of a trade-off between longevity, copper resistance, and fecundity. On the other hand, it is notable that late-physiological age selected females were still laying eggs when exposed to copper media whereas non-selected females were not. Even though the eggs were smaller and presumably of lower quality, the continued oviposition of selected females under copper stress at late physiological age potentially counterbalances this small trade-off. Tests of viability of eggs laid by late-aged females would further clarify this pattern; however, we did not test this. Overall, we found no consistent evidence of a clear cost or trade-off with fecundity in flies with increased longevity and copper resistance under non-stressful conditions.

### Adult selection for copper resistance improved developmental resistance with no costs under control conditions

In addition to reduced fecundity or egg size, reproductive trade-offs have the potential to affect developmental success of future offspring. We therefore examined the developmental success of eggs laid by young females with or without prior selection for adult copper resistance. Development viability (the proportion of 50 eggs that successfully eclosed) was influenced by copper treatment in a dose-dependent manner (F_(3,16)_ = 47.35, P < 0.0001). Compared to control conditions, viability decreased by 13.7% and 62.3% when eggs developed on 5mM and 7mM CuSO_4_ respectively (0mM vs 5mM: adj. P < 0.008; 0mM vs 7mM: adj. P < 0.0001) (**Figure 3**A). Selection history did not directly affect viability (F_(1,16)_ = 0.02, P = 0.89), suggesting that there is no cost of selection for offspring, especially under control conditions. However, we observed a significant interaction between copper treatment and selection history (F_(3,16)_ = 6.58, P < 0.005), which was driven by higher viability of the offspring of copper-selected females under 5mM and 7mM copper stress (selected vs non-selected on 5mM: adj. P < 0.03; selected vs non-selected on 7mM: adj. P < 0.0005; **Figure 3**A).

**Figure 3.**
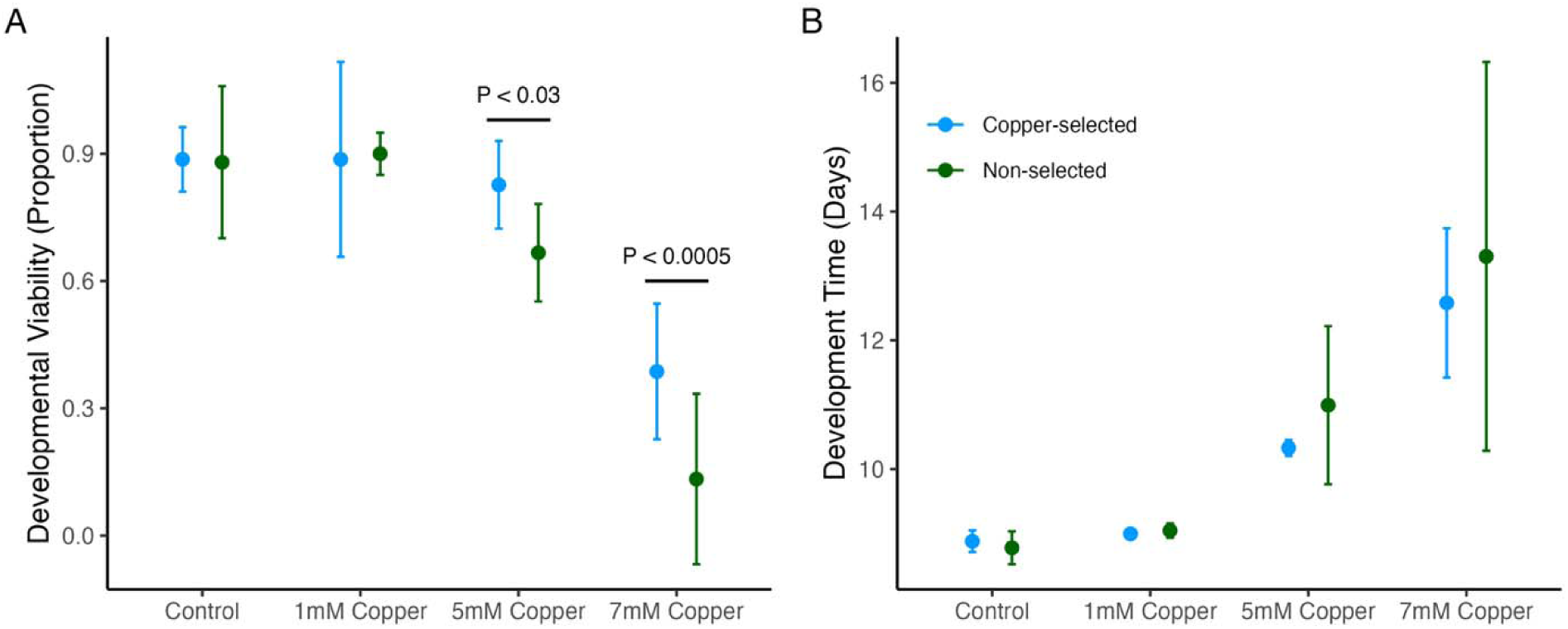
Development success was influenced by copper exposure and selection. A. The proportion of 50 eggs that eclosed declined with increasing copper concentration, and offspring of females that had experienced selection for copper resistance had higher viability when exposed to 5 or 7mM copper (F_(3,16)_ = 6.58, P < 0.005). B. Development time was extended by 1.8-4.1 days due to 5mM and 7mM copper exposure (F_(3,16)_ = 44.26, P < 0.0001). Selection history did not affect development time other than by reducing variability in the response on 5mM and 7mM copper media (F_(1,16)_ = 0.09, P = 0.77). In both plots, offspring of flies from the control population are shown in green; offspring of copper selected flies are shown in blue. Points indicate average estimates +/− 95% CI. Indicated p values were corrected for post-hoc comparisons using the Bonferroni method. **Alt text:** Graphs show differences in developmental viability and development time in two subfigures A and B. Data illustrate average responses due to differences in selection history and experimental treatment of the offspring produced from long-lived and control flies.

Development time was significantly affected by copper treatment but selection history had no direct or interaction effects (Copper Treatment: F_(3,16)_ = 44.26, P < 0.0001; Selection History: F_(1,16)_ = 0.09, P = 0.77; **Figure 3**B). Relative to control conditions, exposure to 5mM or 7mM copper increased development time by 1.8 or 4.1 days, respectively. Variability in the time required to complete development also notably increased for control (non-selected) flies, with some individuals requiring nearly 7 additional days when exposed to 7mM copper (**Figure 3**B).

Sex ratio (males to females) was not significantly affected by copper treatment or by selection history (Copper Treatment: F_(3,16)_ = 0.05, P = 0.98; Selection History: F_(1,16)_ = 0.07, P = 0.80), suggesting that the effect of copper exposure on male and female development success was similar regardless of concentration. Overall, we found no evidence of a cost associated with developmental success of offspring from females that had experienced selection for copper resistance, suggesting that the fitness benefits associated with increased longevity are not counterbalanced by lower reproductive output, quality, or success.

## Discussion

In a recent study, long-term artificial selection for copper resistance resulted in an unexpected, correlated increase in longevity (Arnold and Everman 2026). In that same study, flies that were not exposed to copper selection lost resistance, suggesting that maintenance of resistance to copper stress is costly. A standard paradigm in evolutionary biology proposes that organisms do not have infinite energetic resources at their disposal and that enhancement or optimization of one energetically demanding trait could reasonably come at a cost to other energetically demanding traits (Stephen 1992; Zera and Harshman 2001). The response to copper toxicity has been shown to be energetically costly in gastropods (Osborne and Prosser 2025b), and energetic trade-offs between lifespan and fecundity under the disposable soma theory (Kirkwood 1977) have long been debated (Rose 1984; Flatt 2011; Cohen et al. 2020). The correlated response between increased copper resistance and lifespan observed in copper-selected flies argues against a resistance-longevity tradeoff to account for the loss of copper resistance in control populations. In this study, we explored potential trade-offs between copper resistance and reproduction with the goal of illuminating trade-offs that may exist between copper resistance and other aspects of life history.

Surprisingly, we found no evidence of a clear trade-off between copper resistance, longevity, and reproduction. In fact, under control experimental conditions copper selected females out-lived non-selected females (**Figure 1**), laid more eggs (**Figure 2**), and produced offspring with similar viability and development time compared with non-selected females (**Figure 3**). Even if the slight increase in lifetime fecundity is dismissed given the minor effect, we still found no evidence of a copper resistance cost in the reproductive traits assessed. Under copper stress conditions, copper selection history had no statistically significant effect on the metrics of fecundity we assessed (age-specific fecundity, total fecundity, and egg quality) but copper selection did improve viability of offspring under moderate copper stress relative to the offspring of non-selected females (**Figure 3**). Increased offspring survival from flies that experienced copper selection during the adult life state only was not necessarily expected; we previously found evidence of potential genetic decoupling of life stage-specific resistance to copper stress (Everman et al. 2021) and decoupling of stress traits across developmental stages has been reported for other traits in *D. melanogaster* (Freda et al. 2017, 2019). Our findings demonstrate that copper-resistance that evolved through artificial selection during the adult life stage only can influence resistance during development.

To our knowledge, our results are contradictory to the only other study to examine fitness costs of metal adaptation in artificially selected *D. melanogaster*. Shirley and Sibly (1999) performed an artificial selection experiment for 20 generations to increase cadmium resistance. Similar to our study, they succeeded in generating cadmium-resistant flies that outperformed non-selected flies under cadmium stress conditions. However, when assessed under control conditions, fecundity of cadmium-resistant flies was reduced by 44% compared to non-selected flies, suggesting a reproductive cost of increased cadmium resistance (Shirley and Sibly 1999). Unlike the essential micronutrient copper, cadmium is a non-essential heavy metal that is considered dangerous at very low doses (Charkiewicz et al. 2023). Despite differences in micronutrient requirement and dose-dependent toxicity, cadmium and copper have similar reactivity, size, and oxidative stress effects at the cellular level (Ercal et al. 2001; Belyaeva et al. 2012; Husak 2015). Additionally, the copper-selected flies used in this study retained their starting levels of cadmium resistance throughout the selection period in contrast to non-selected flies which lost cadmium resistance (Arnold and Everman 2026). The lack of complete independence between copper and cadmium resistance led us to expect a similar reproductive cost of copper resistance. However, the reproductive cost that accompanied cadmium resistance in Shirley and Sibly (1999) may be unique to adaptation to cadmium stress (see also Sadeq and Beckerman 2019), or may be the result of unique, polygenic responses to selection that have case-by-case consequences for fitness traits.

Although we could not link the costs of maintaining copper resistance to trade-offs with classic lifespan and reproductive fitness traits, multiple alternative avenues remain to be explored. First, body condition, scope for growth, respiration, and metabolism may differ between the copper-selected and non-selected flies used in our study. For example, adaptation of the worm *Nereis diversicolor* to high soil levels of zinc and copper was associated with decreased growth rate, body condition (lipid and carbohydrate content of tissues), and altered metabolic rate (Pook et al. 2009). Assessment of energy budgets of copper-selected and non-selected flies may reveal metabolic costs of resistance. Second, lifespan and reproductive fitness costs of copper resistance may be context-dependent (Lenormand et al. 2018). We assessed the effects of a low dose of copper on fecundity to avoid strong mortality effects on study females from the selected and non-selected populations. However, at more stressful doses, or under different copper exposure conditions (such as during maternal development), life history costs may be revealed.

Adaptation of populations to high levels of metal contamination has been reported in microbes (Adamo et al. 2012; Bazzicalupo et al. 2025), plants (Courbot et al. 2007), invertebrates (e.g. Lopes et al. 2004; Pook et al. 2009; Green et al. 2022; Arnold and Everman 2026), and vertebrates (Blanchette et al. 2025). However, the consequences for the persistence of these metal-adapted populations from the perspective of fitness trade-offs is less often examined. Artificial selection experiments in model organisms such as *D. melanogaster* provide insight into the complex effects of adaptive evolution to chemical stress. Our study consistently points to a cost of maintaining heavy metal resistance, but in our study populations, metal resistance does not come at a cost to lifespan or reproduction.

## Data Availability

All data are available from FigShare upon acceptance

## Author Contributions

CMR, KAA, and ERE collected data, ERE and CMR performed the analysis and constructed plots, CMR wrote the original draft. ERE conceptualized and supervised the project, developed methodology, curated data, revised and edited the manuscript, and acquired funding for the project.

## Acknowledgments

We thank Charlton Miller for assistance with ImageJ and Paul Klawinski for advice on statistical analysis.

## References

Abràmoff, M. D., P. J. Magalhães, and S. J. Ram. 2004. Image procesing with ImageJ. Biophotonics Int. 11:36–42.

Adamo, G. M., S. Brocca, S. Passolunghi, B. Salvato, and M. Lotti. 2012. Laboratory evolution of copper tolerant yeast strains. Microb. Cell Factories 11:1.

Agra, A. R., A. M. V. M. Soares, and C. Barata. 2011. Life-history consequences of adaptation to pollution. “*Daphnia longispina* clones historically exposed to copper.” Ecotoxicology 20:552–562.

Arnold, K. A., and E. R. Everman. 2026. Artificial selection for resistance to copper and off-target physiological and behavioral effects in *Drosophila melanogaster*. Ecotoxicol. Environ. Saf. 312:119974.

Backhaus, B., E. Sulkowski, and F. W. Schlote. 1984. A semi-synthetic, general-purpose medium for *Drosophila melanogaster*. Drosoph. Inf. Serv. 60:210–212.

Bazzicalupo, A. L., P. C. Kahn, E. Ao, J. Campbell, and S. P. Otto. 2025. Evolution of cross-tolerance to metals in yeast. PNAS 122.

Bazzicalupo, A. L., J. Ruytinx, Y.-H. Ke, L. Coninx, J. V. Colpaert, N. H. Nguyen, R. Vilgalys, and S. Branco. 2019. Incipient local adaptation in a fungus: evolution of heavy metal tolerance through allelic and copy-number variation. bioRxiv, doi: 10.1101/832089.

Blanchette, A., K.-J. Su, A. J. Frick, J. Karubian, and A. R. Gunderson. 2025. Unprecedented lead tolerance in an urban lizard. Environ. Res. 285:122531.

Calap-Quintana, P., J. González-Fernández, N. Sebastiá-Ortega, J. Llorens, and M. Moltó. 2017. *Drosophila melanogaster* models of metal-related human diseases and metal toxicity. Int. J. Mol. Sci. 18:1456.

Charkiewicz, A. E., W. J. Omeljaniuk, K. Nowak, M. Garley, and J. Nikliński. 2023. Cadmium toxicity and health effects—a brief summary. Molecules 28:6620.

Cohen, A. A., C. F. D. Coste, X. Li, S. Bourg, and S. Pavard. 2020. Are trade-offs really the key drivers of ageing and life span? Funct. Ecol. 34:153–166.

Courbot, M., G. Willems, P. Motte, S. Arvidsson, N. Roosens, P. Saumitou-Laprade, and N. Verbruggen. 2007. A major quantitative trait locus for cadmium tolerance in *Arabidopsis halleri* colocalizes with *HMA4*, a gene encoding a heavy metal ATPase. Plant Physiol. 144:1052–1065.

Das, D., M. Moniruzzaman, A. Sarbajna, and S. B. Chakraborty. 2017. Effect of heavy metals on tissue-specific antioxidant response in Indian major carps. Environ. Sci. Pollut. Res. 24:18010–18024.

Eränen, J. K. 2008. Rapid evolution towards heavy metal resistance by mountain birch around two subarctic copper–nickel smelters. J. Evol. Biol. 21:492–501.

Everman, E. R., K. M. Cloud-Richardson, and S. J. Macdonald. 2021. Characterizing the genetic basis of copper toxicity in *Drosophila* reveals a complex pattern of allelic, regulatory, and behavioral variation. Genetics 217:1–20.

Everman, E. R., S. J. Macdonald, and J. K. Kelly. 2023. The genetic basis of adaptation to copper pollution in *Drosophila melanogaster*. Front. Genet. 14:1144221.

Everman, E. R., C. L. McNeil, J. L. Hackett, C. L. Bain, and S. J. Macdonald. 2019. Dissection of complex, fitness-related traits in multiple *Drosophila* mapping populations offers insight into the genetic control of stress resistance. Genetics 211:19.

ffrench-Constant, R. H., and C. Bass. 2017. Does resistance really carry a fitness cost? Curr. Opin. Insect Sci. 21:39–46.

Fisker, K. V., J. G. Sørensen, C. Damgaard, K. L. Pedersen, and M. Holmstrup. 2011. Genetic adaptation of earthworms to copper pollution: is adaptation associated with fitness costs in *Dendrobaena octaedra?* Ecotoxicology 20:563–573.

Flatt, T. 2011. Survival costs of reproduction in *Drosophila*. Exp. Gerontol. 46:369–375.

Fox, J., S. Weisberg, and B. Price. 2001. car: Companion to Applied Regression.

Freda, P. J., J. T. Alex, T. J. Morgan, and G. J. Ragland. 2017. Genetic decoupling of thermal hardiness across metamorphosis in *Drosophila melanogaster*. Integr. Comp. Biol. 57:999–1009.

Freda, P. J., Z. M. Ali, N. Heter, G. J. Ragland, and T. J. Morgan. 2019. Stage-specific genotype-by-environment interactions for cold and heat hardiness in *Drosophila melanogaster*. Heredity 123:479–491.

Garland, T., C. J. Downs, and A. R. Ives. 2022. Trade-offs (and constraints) in organismal biology. Physiol. Biochem. Zool. 95:82–112.

Georgieva, S. S., S. P. McGrath, D. J. Hooper, and B. S. Chambers. 2002. Nematode communities under stress: the long-term effects of heavy metals in soil treated with sewage sludge. Appl. Soil Ecol. 20:27–42.

Gerstein, A. C., J. Ono, D. S. Lo, M. L. Campbell, A. Kuzmin, and S. P. Otto. 2015. Too much of a good thing: The unique and repeated paths toward copper adaptation. Genetics 199:555–571.

Green, L., M. Coronado-Zamora, S. Radío, G. E. Rech, J. Salces-Ortiz, and J. González. 2022. The genomic basis of copper tolerance in *Drosophila* is shaped by a complex interplay of regulatory and environmental factors. BMC Biol. 20:275.

Harshman, L. G., K. M. Moore, M. A. Sty, and M. M. Magwire. 1999. Stress resistance and longevity in selected *lines of Drosophila melanogaster*. Neurobiol. Aging 20:521–529.

Hope. 2012. Rmisc: Ryan Miscellaneous.

Husak, V. 2015. Copper and copper-containing pesticides: metabolism, toxicity and oxidative stress. J. Vasyl Stefanyk Precarpathian Natl. Univ. 2:38–50.

Jomova, K., D. Vondrakova, M. Lawson, and M. Valko. 2010. Metals, oxidative stress and neurodegenerative disorders. Mol. Cell. Biochem. 345:91–104.

Kirkwood, T. B. L. 1977. Evolution of aging. Nature 270:301–304.

Küpper, H., E. Lombi, F. Zhao, G. Wieshammer, and S. P. McGrath. 2001. Cellular compartmentation of nickel in the hyperaccumulators *Alyssum lesbiacum*, *Alyssum bertolonii* and *Thlaspi goesingense*. J. Exp. Bot. 52:2291–2300.

Lee, K. P., S. J. Simpson, F. J. Clissold, R. Brooks, J. W. O. Ballard, P. W. Taylor, N. Soran, and D. Raubenheimer. 2008. Lifespan and reproduction in *Drosophila*: New insights from nutritional geometry. Proc. Natl. Acad. Sci. 105:2498–2503.

Lenormand, T., N. Harmand, and R. Gallet. 2018. Cost of resistance: an unreasonably expensive concept. Rethink. Ecol. 3:51–70.

Lenth, R. 2025. emmeans: Estimated Marginal Means, aka Least-Squared Means.

Lopes, I., D. J. Baird, and R. Ribeiro. 2004. Genetic determination of tolerance to lethal and sublethal copper concentrations in field populations of *Daphnia longispina*. Arch. Environ. Contam. Toxicol. 46:43–51.

Maestri, E., M. Marmiroli, G. Visioli, and N. Marmiroli. 2010. Metal tolerance and hyperaccumulation: Costs and trade-offs between traits and environment. Environ. Exp. Bot. 68:1–13.

Osborne, R. K., and R. S. Prosser. 2025a. Assessing the energetic cost of exposure to copper in a freshwater gastropod. Environ. Sci. Pollut. Res. 32:18820–18831.

Osborne, R. K., and R. S. Prosser. 2025b. Assessing the energetic cost of exposure to copper in a freshwater gastropod. Environ. Sci. Pollut. Res. 32:18820–18831.

Papadopulos, A. S. T., A. J. Helmstetter, O. G. Osborne, A. A. Comeault, D. P. Wood, E. A. Straw, L. Mason, M. F. Fay, J. Parker, L. T. Dunning, A. D. Foote, R. J. Smith, and J. Lighten. 2021. Rapid parallel adaptation to anthropogenic heavy metal pollution. Mol. Biol. Evol. 38:3724–3736.

Pook, C., C. Lewis, and T. Galloway. 2009. The metabolic and fitness costs associated with metal resistance in *Nereis diversicolor*. Mar. Pollut. Bull. 58:1063–1071.

R Core Team. 2023. R: A language and environment for statistical computing. R Foundation for Statistical Computing, Vienna, Austria.

Rasband, W. S. 2007. ImageJ. National Institutes of Health, Bethesda, Maryland.

Rivero, A., A. Magaud, A. Nicot, and J. Vézilier. 2011. Energetic cost of insecticide resistance in *Culex pipiens* mosquitoes. J. Med. Entomol. 48:694–700.

Rose, M. R. 1984. Laboratory evolution of postponed senescence in *Drosophila melanogaster*. Evolution 38:1004.

Rose, M. R., L. N. Vu, S. U. Park, and J. L. Graves, Jr. 1992. Selection on stress resistance increases longevity in *Drosophila melanogaster*. Exp. Gerontol. 27:241–250.

Ruiz, L. M., A. Libedinsky, and A. A. Elorza. 2021. Role of copper on mitochondrial function and metabolism. Front. Mol. Biosci. 8:711227.

Sadeq, S. A., and A. P. Beckerman. 2019. The chronic effects of copper and cadmium on life history traits across Cladocera species: A meta-analysis. Arch. Environ. Contam. Toxicol. 76:1–16.

Service, P. M., E. W. Hutchinson, M. D. MacKinley, and M. R. Rose. 1985. Resistance to environmental stress in *Drosophila melanogaster* selected for postponed senescence. Physiol. Zool. 58:380–389.

Shirley, M. D. F., and R. M. Sibly. 1999. Genetic basis of a between-environment trade-off involving resistance to cadmium in *Drosophila melanogaster*. Evolution 53:826–836.

Stephen, S. 1992. The Evolution of Life Histories. Oxford University Press, New York, NY.

Wickham, H., M. Averick, J. Bryan, W. Chang, L. McGowan, R. François, G. Grolemund, A. Hayes, L. Henry, J. Hester, M. Kuhn, T. Pedersen, E. Miller, S. Bache, K. Müller, J. Ooms, D. Robinson, D. Seidel, V. Spinu, K. Takahashi, D. Vaughan, C. Wilke, K. Woo, and H. Yutani. 2019. Welcome to the Tidyverse. J. Open Source Softw. 4:1686.

Zera, A. J., and L. G. Harshman. 2001. The physiology of life history trade-offs in animals. Annu. Rev. Ecol. Syst. 32:95–126.

Zwaan, B., R. Bijlsma, and R. F. Hoekstra. 1995. Direct selection on life span in *Drosophila melanogaster*. Evolution 49:649–659.

